# Inter-joint and temporal coordination change in very slow walking

**DOI:** 10.1101/588095

**Authors:** Virginia L. Little, Theresa E. McGuirk, Carolynn Patten

## Abstract

**Purpose:** Very slow walking has been suggested to be a distinctively different motor behavior than walking at comfortable gait speeds. While kinematic and spatiotemporal gait parameters are known to scale with gait speed, inter-joint coordination during swing remains consistent, at least across comfortable speeds. The purpose of this study was to determine whether coordination patterns serving limb clearance and shortening differ with very slow walking, providing additional support for the premise that very slow walking represents a unique motor behavior.

**Methods:** We assessed nine healthy adults walking overground at their self-selected speed and two-to-three progressively slower speeds. We collected lower extremity kinematics with 3D motion analysis and quantified joint motion contributions to limb clearance and shortening. We investigated changes in coordination using linear mixed models to determine magnitude and timing differences of joint influence across walking speeds.

**Results:** Hip and knee influences serving limb clearance reduced considerably with slower walking speeds. Similarly, knee influence on limb shortening reduced with very slow walking. Importantly, ankle influence remained unchanged across gait speeds for limb shortening and reduced subtly for limb clearance. Temporally, joint influences on limb clearance varied across walking speeds. Specifically, the temporal order of peak hip and knee influences reversed between comfortable and very slow walking. For limb shortening the timing of ankle influence remained unchanged while the timing of knee influence occurred later in the gait cycle for slower walking speeds.

**Conclusions:** Our results demonstrate temporal coordination and the relative joint contributions serving limb clearance and shortening differ with very slow walking providing additional evidence that slow walking may be a behavior distinct from walking at comfortable speeds.

## Introduction

Humans can walk at a variety of speeds and typically modulate their gait based on environment and task demands.(Warren 2018; Kesler et al. 2005; Sun et al. 1996; Licence et al. 2015) For instance, individuals typically adopt a slower gait speed in the dark, on sloped surfaces, and in crowds.(Kesler et al. 2005; Sun et al. 1996) Similarly, concurrent competing task goals, such as increased cognitive load or obstacle negotiation, often coincide with slowed gait.(Licence et al. 2015) In extreme cases, for instance when walking with family members or friends who have sustained an injury, an individual might notice walking becomes halting and lacks the rhythm and automaticity characteristic of natural bipedal walking. This phenomenon begs the question of whether walking at very slow speeds is inherently different than walking at usual or comfortable gait speed.

Understanding unique characteristics of very slow gait offers scientific importance but also has implications for gait rehabilitation. Gait researchers commonly speed-match, using healthy individuals walking at very slow speeds, for a direct comparison with pathologic gait.(Lehmann et al. 1987; Chen et al. 2005; Little, McGuirk, and Patten 2014) However, it has been suggested that very slow walking (<0.4-0.5 m/sec) may be a unique motor behavior utilizing a different control strategy than walking at comfortable walking speeds.(Leiper and Craik 1991; Smith and Lemaire 2018) If this were the case, speed-matching for biomechanical studies may not always be appropriate. Furthermore, if very slow walking is a unique behavior, then contrary to common thought, individuals working to recover walking ability after neurologic injury, such as a stroke or spinal cord injury, may be confronted with learning a new motor behavior rather than re-learning a well-practiced motor task. For these reasons, understanding very slow gait becomes important.

Kinematic and spatiotemporal features of gait are known to scale with gait speed, but inter-joint coordination during swing is thought to remain consistent across gait speeds.(Kirtley, Whittle, and Jefferson 1985; Nymark et al. 2005; Oberg, Karsznia, and Oberg 1994; Stoquart, Detrembleur, and Lejeune 2008; Shemmell et al. 2007; Mentiplay et al. 2018) However, studies investigating the relationship of gait characteristics to gait speed typically do not include very slow speeds, limiting potential insight regarding this behavior.(Kirtley, Whittle, and Jefferson 1985; Oberg, Karsznia, and Oberg 1994; Shemmell et al. 2007) Indeed, when spatiotemporal characteristics of gait have been investigated at very slow gait speeds, the relationship between temporal characteristics and gait speed have been found to differ between very slow and comfortable gait speeds.(Smith and Lemaire 2018) It seems reasonable to posit that coordination at very slow gait speeds may also differ from the coordination used at typical or comfortable walking speeds. However, this question remains to be systematically investigated and is the purpose of our study.

We are specifically interested in the inter-joint and temporal coordination patterns involved in limb shortening and limb clearance during the swing phase of gait. Coordinated motion of the stance and swing limbs serve the objective of repositioning the swing limb from behind to in front of the stance limb with sufficient clearance to avoid premature foot contact.(Gage 1990; Perry 1992) For this investigation, we focused our study on the timing and magnitude of contributions to limb shortening and limb clearance directly attributable to the sagittal plane joint motions of the swing limb. To determine whether coordination patterns change with very slow walking, we assessed the joint contributions across a range of walking speeds spanning gait speeds consistent with comfortable walking in health to very slow walking.

## Methods

### Participants

We studied 9 healthy adults (age: 43.7 ± 11.2 yrs; men/women: 5/4) free from any cardiac, orthopedic, or neurologic conditions that would limit their ability to walk. All participants provided written informed consent approved by the Stanford University Institutional Review Board prior to enrollment.

### Data Collection and Processing

We studied participants while they walked overground at their self-selected speed (SSWS) and up to three progressively slower speeds. Rather than utilizing external constraints (e.g., timing gate, metronome), we instructed the participants to ‘walk slower’, ‘walk even slower’, and if possible, ‘walk even slower’. This allowed us to investigate how participants self-organized slower walking behavior and the relationship between gait speed and joint influence on limb clearance and limb shortening.

All participants wore their own footwear, typically a flat, athletic style shoe, and their walking was not constrained by external pacing. We obtained and labeled three-dimensional marker data using a seven-camera motion capture system (Qualisys AB., Gothenburg, Sweden, 100Hz) and a modified Cleveland Clinic marker set (5 clusters and 23 additional markers) as described by Chen & Patten.(Chen and Patten 2008) In Visual 3D Basic (v 3.99.25.7, C-Motion, Germantown, MD), we modeled and filtered (lowpass 4^th^ order Butterworth, 6 Hz cutoff) marker data and calculated kinematics. We time-normalized kinematic data to a 101-point gait cycle using custom Matlab (MathWorks Version 7.7.0 R2008b, Natick, MA) scripts.

### Biomechanical Model

We used a planar model of the leg to investigate the relative contributions of sagittal plane swing limb joint angles to limb clearance and limb shortening.(Moosabhoy and Gard 2006) By convention the model reports hip flexion, knee flexion, and ankle dorsiflexion angles as positive joint rotations. The model used in the current analysis was developed by Moosabhoy and Gard and is described briefly below.(Moosabhoy and Gard 2006)

#### Limb clearance sensitivity

Toe clearance served as our proxy for limb clearance. We used the vertical trajectory of the distal toe marker to quantify toe position. Vertical toe position (i.e., toe height) is a function of: i) vertical hip position, ii) thigh, shank, and foot segment lengths, and iii) hip, knee, and ankle angles (*see Equation 2* (Moosabhoy and Gard 2006)). As such, the relative contribution of each joint, or sensitivity, can be determined by calculating the partial derivative of the vertical toe position with respect to each joint angle (*see Equations 6-8* (Moosabhoy and Gard 2006)). Interpretation of these values is based on the instantaneous direction of joint motion.

#### Limb shortening sensitivity

Shortening of the swing limb, rather than an absolute measure of clearance, provides a direct measure of the capacity for limb shortening to enable the swing limb to advance in front of the body without foot-floor contact.(Moosabhoy and Gard 2006) We quantified limb shortening as the percent reduction in normalized limb length relative to the instantaneous height of the hip joint center. Normalized limb length was calculated as the instantaneous hip-toe distance (HTD) divided by the instantaneous vertical distance from the hip joint center to the floor (HFD).(Moosabhoy and Gard 2006) The hip-toe distance is calculated via the Pythagorean theorem using the vertical and fore-aft coordinates of the hip joint center and distal toe marker (*see Equation 15* (Moosabhoy and Gard 2006)). Normalized limb length values less than 1 indicate limb shortening. Again, the partial derivative of normalized limb length with respect to the contributing joints (i.e., knee and ankle) quantifies the relative contribution of the knee and ankle to limb shortening (*see Equations 16-17* (Moosabhoy and Gard 2006)).

### Outcomes

#### Inter-joint coordination

Interpretation of limb clearance and limb shortening sensitivity values requires simultaneous knowledge of the direction of joint motion. For example, at mid-swing, limb clearance sensitivity with respect to knee flexion is positive, suggesting knee flexion increases limb clearance. However, the knee is extending during this time, thus motion at the knee reduces limb clearance around mid-swing (~80% of gait cycle; Figure 1). Accordingly, we quantified the estimated *joint influence* on limb clearance (LCI) and limb shortening (LSI) throughout the cycle using the following equation:

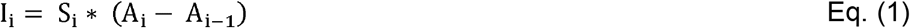

where I_i_ is the influence value, S_i_ is the sensitivity value, and A_i_ is the sagittal plane joint angle at time i. It follows that positive influence values for limb clearance indicate the specified joint increases limb clearance. For limb shortening a negative influence value indicates shortening of the limb by the given joint.

**Fig 1.**
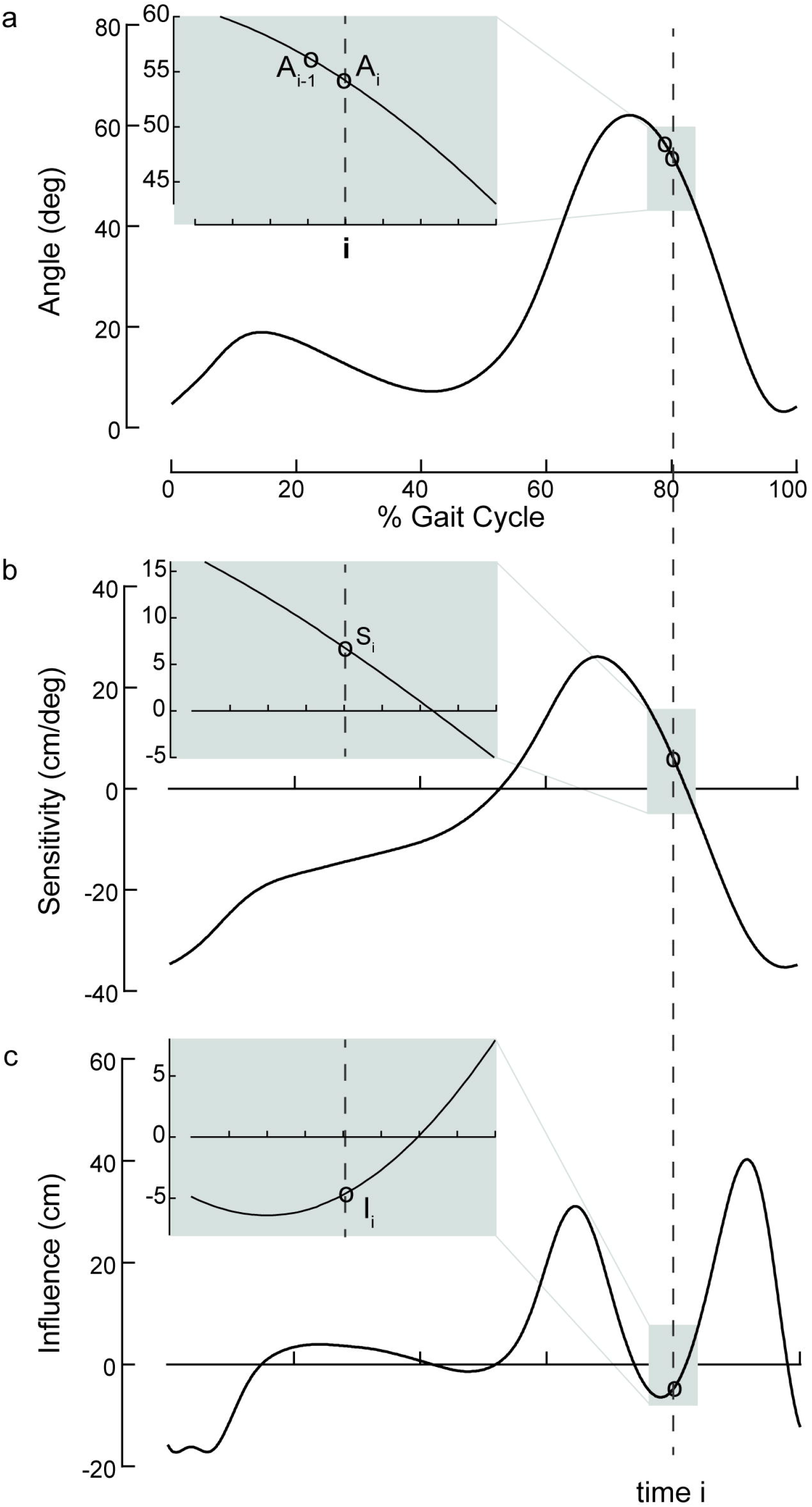
Quantification of joint influence. The relationship between: a) sagittal plane joint angle, b) limb clearance sensitivity, and c) limb clearance influence, illustrated using group mean data at self-selected walking speed. Estimated *joint influence* on limb clearance (LCI) and limb shortening (LSI) throughout the gait cycle are quantified using the following general equation: I_i_ = S_i_ * (A_i_ − A_i−1_); where I_i_ is the influence value, S_i_ is the sensitivity value, and A_i_ is the sagittal plane joint angle, all at time i represented by the dashed vertical line throughout all subplots. Shaded grey regions at time i highlight individual equation components; these are expanded in insets to illustrate greater detail between 76-84% of the gait cycle. Incorporation of the direction of joint motion facilitates interpretation of sensitivity values. Specifically, at time i the sensitivity value (subplot b) is positive, suggesting joint motion increases limb clearance. However, because the joint is extending (subplot a), the influence value is negative, indicating the joint motion at time i reduces limb clearance

Previous studies of toe clearance and fall risk in healthy elders investigated toe clearance in mid-swing(Murray and Clarkson 1966; Begg et al. 2007) and late swing.(Mills, Barrett, and Morrison 2008) However, we noted the critical toe clearance, identified by a local minimum of the vertical trajectory of the toe in mid-swing(D. A. Winter 1992; Moosabhoy and Gard 2006; Murray and Clarkson 1966; Begg et al. 2007), is often absent in healthy controls when walking at slow speeds (Figure 2).(Santhiranayagam et al. 2017) Thus, to maintain consistency throughout our sample, all influence values were investigated at peaks relevant to the task goals of limb clearance and limb shortening. For limb clearance (Figure 3a): i) the hip influence peak was negative and typically occurred in the first half of swing (for n=2 the peak occurred just prior to toe off), ii) the knee peak was positive and typically occurred in the first half of swing (for n=2 the peak occurred just prior to toe off), and iii) the ankle peak was positive and occurred around mid-swing. For limb shortening (Figure 4a), both influence peaks investigated were negative and occurred within the first half of swing.

**Fig 2.**
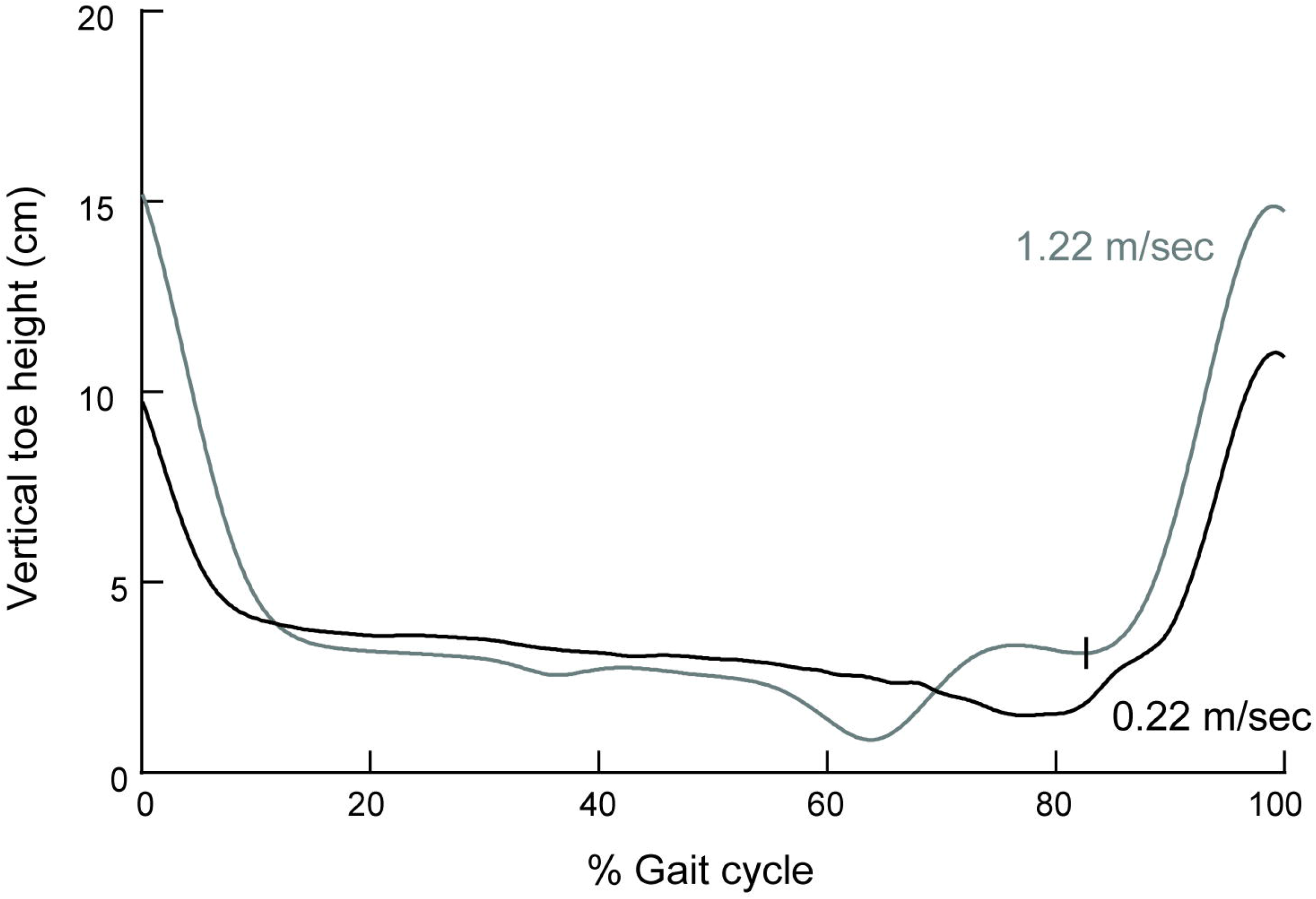
Vertical toe height. The vertical trajectory of the great toe marker, time normalized to the gait cycle, illustrates limb clearance. Data are from a single subject walking at their self-selected walking speed (1.22 m/sec; grey) and a very slow walking speed (0.22 m/sec; black). The local minimum (denoted with vertical hashmark) of the vertical trajectory of the great toe marker, typically used to identify critical toe clearance in mid-swing, is absent at slow walking speeds in some individuals, as illustrated here. As a result, all influence values were investigated at their first peak during swing

**Fig 3.**
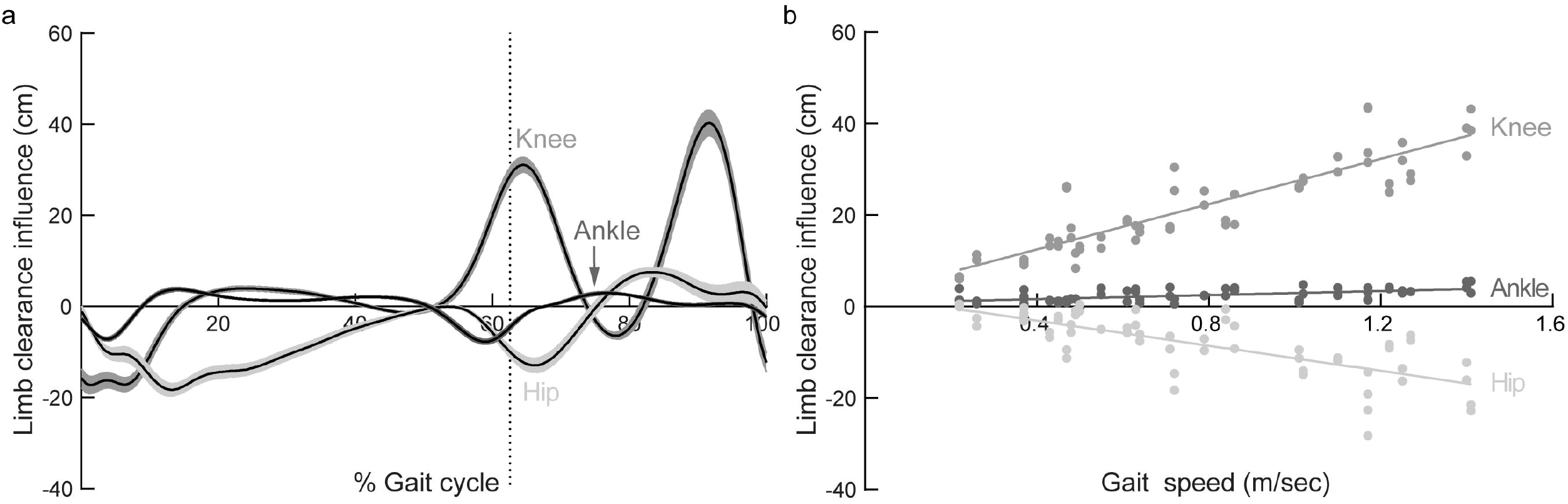
Limb clearance influence a) The general pattern of limb clearance influence, time-normalized to the gait cycle, is depicted with respect to the hip, knee, and ankle. Data are mean ± SEM at self-selected walking speeds. Dotted vertical line represents timing of toe off. The peaks quantified for analysis occur during swing as follows: i) hip (negative peak), ii) knee (1^st^ positive peak), and iii) ankle (positive peak). Positive influence values for limb clearance indicate the specified joint increases limb clearance. b) Peak limb clearance influence represented across gait speeds with notable increases and decreases in peak hip and knee influence, respectively, as speed is reduced

**Fig 4.**
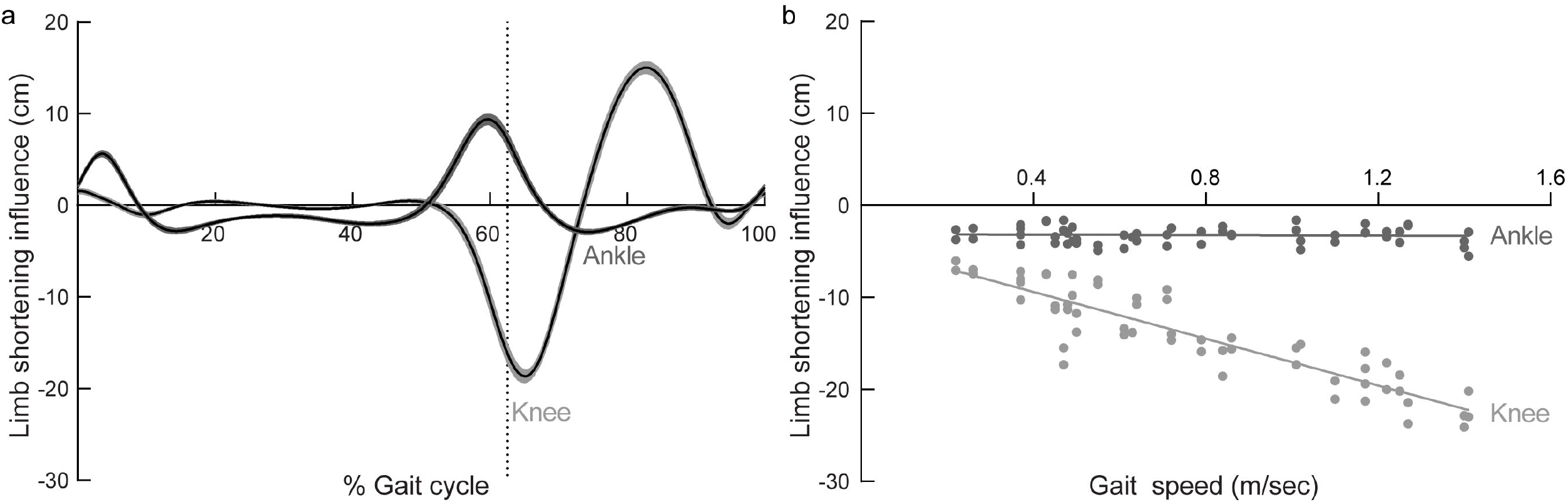
Limb shortening influence a) The general pattern of limb shortening influence, time-normalized to the gait cycle, is depicted with respect to the knee and ankle. Data are mean ± SEM at self-selected walking speeds. Dotted vertical line represents timing of toe off. The peaks quantified for analysis occur during swing as follows: i) knee (negative peak), and ii) ankle (negative peak). Positive influence values for limb shortening indicate lengthening of the limb by the given joint. b) Peak limb shortening influence represented across gait speeds with systematic decreases in knee influence as speed is reduced

We investigated the joint influences of limb clearance and limb shortening as they relate to gait speed. Understanding these concurrent relationships allows us to describe how the inter-joint coordination, or relative joint contributions to limb clearance and limb shortening, may differ at very slow walking speeds.

#### Temporal coordination

To assess the temporal coordination pattern between joints, we also investigated the timing of peak influence for each joint, relative to the gait cycle. We identified altered temporal coordination in a manner consistent with inter-joint coordination.

#### Proportional influence

To describe the relative contributions from each joint, we quantified the proportional influence of each joint serving limb clearance and limb shortening. To do this, we summed (absolute) discrete values (i.e., all values made positive) of influence magnitude at the instant of peak influence for each joint and calculated the proportion of influence ascribed to each joint for a given task.

### Statistical Analysis

We pooled the data from both legs for all analyses. We used descriptive statistics to quantify proportional influences of each joint. For all remaining variables, we used Goodness of Fit to test for normality and Levene’s test to assess equality of variances. Minor violations were noted for the parametric assumptions of normality and homogeneity of variances (p’s > 0.05).

To accommodate these violations, we used linear mixed models to determine if the fixed effects of gait speed, joint, and the interaction between gait speed and joint were significant predictors of peak limb clearance and limb shortening influences.(Quené and van den Bergh 2004) In each of these models we included random effects including intercepts for each joint and by-joint random slopes for the effect of gait speed.(B. Winter 2013) Finally, we used the Unequal Variance covariance structure with multiple repeats for each subject. To understand the non-linear trend apparent in our timing variables, we assessed the appropriateness of a quadratic term (gait speed x gait speed) in the linear mixed models used to identify predictors of the timing of peak influences with respect to (wrt) the gait cycle.

As an exploratory analysis to determine if the temporal order of hip and knee influences on limb clearance shifted across gait speeds, we calculated the difference between knee and hip influence timing relative to the gait cycle. Positive values of timing difference indicate the knee influence occurs later in the gait cycle than the hip influence. We used a linear mixed model with a fixed effect of gait speed, allowing for varying intercepts by subject, and the Unequal Variance covariance structure with multiple repeats for each subject to determine the relationship between gait speed and the difference between knee and hip influence timings.

In total, we performed 5 linear mixed models; after correcting for multiple comparisons, we established statistical significance at: α = 0.01 for all variables. We then conducted separate models for each joint removing the main effect of joint and the interaction term involving joint to investigate unique relationships by joint. The Type I error rate was carried through and used for all subsequent models. All statistical tests were performed with JMP^®^ Pro 14.0.0 (SAS Institute Inc., Cary, NC).

## Results

Our participants produced a range of walking speeds: 0.22 – 1.41 m/sec.

### Inter-joint coordination

#### Limb clearance

Across speeds, the knee remained the predominant contributor to limb clearance, consistently providing over half (56-95%) of the influence on limb clearance. At typical walking speeds, the primary swing limb joint contributions to limb clearance are characterized by a dynamic interplay between hip and knee flexion; while knee flexion is the primary contributor to limb clearance, hip flexion counters this objective when the limb is posterior to the trunk in early swing (Figure 3a). Our overall model confirmed that the fixed effects of joint (F_(2,22.9)_=214.8; *p*<0.0001), gait speed (F_(1,19.2)_=23.6; *p*=0.0001), and the interaction between joint and gait speed (F_(2,18.3)_=144.1; *p*<0.0001) were significant predictors of limb clearance influences. Individual joint models revealed the fixed effect of gait speed to be a consistently significant predictor of limb clearance influence. While these findings suggest the influence of sagittal plane joint motion varies across gait speeds, these relationships differ by joint. The relationship of the knee influence on limb clearance reveals a large positive slope (*b*_knee_=24.89; *p*<0.0001); whereas the relationship of the hip influence on limb clearance is characterized by a moderate negative slope (*b*_hip_=−13.76; *p*<0.0001). While the changes noted at the hip and knee across gait speeds are considerable, the relationship between ankle influence on limb clearance and gait speed is subtle (*b*_ankle_=2.28; *p*=0.0005; Figure 3b).

#### Limb shortening

The knee was also the predominant contributor to limb shortening, consistently providing 62-92% of the influence across speeds. In contrast, the ankle contributed only 8-38% of influence on limb shortening (Figure 4a). Our overall model confirmed that the fixed effects of joint (F_(1,10.4)_=394.7; *p*<0.0001), gait speed (F_(1,53.9)_=348.1; *p*<0.0001), and the interaction between joint and gait speed(F_(1,53.9)_=351.0; *p*<0.0001) were significant predictors of limb shortening influences. Individual joint models revealed the fixed effect of gait speed was a significant predictor of limb shortening influence from the knee, but not the ankle. These findings suggest the influence of sagittal plane knee motion varies across gait speeds characterized by a moderate negative slope (*b*_knee_=−12.7; *p*<0.0001). Yet, the influence from the ankle remains unchanged regardless of gait speed (*p*=0.74; Figure 4b).

### Temporal coordination wrt the gait cycle

For limb clearance, the ankle influence occurs later (73-89%) in the gait cycle than the knee (61-82%) and hip (62-80%) influences (Figures 3a and 5a). A similar pattern for the timing of ankle (72-84%) and knee (62-82%) influences was noted for limb shortening (Figures 4a and 5c).

**Fig 5.**
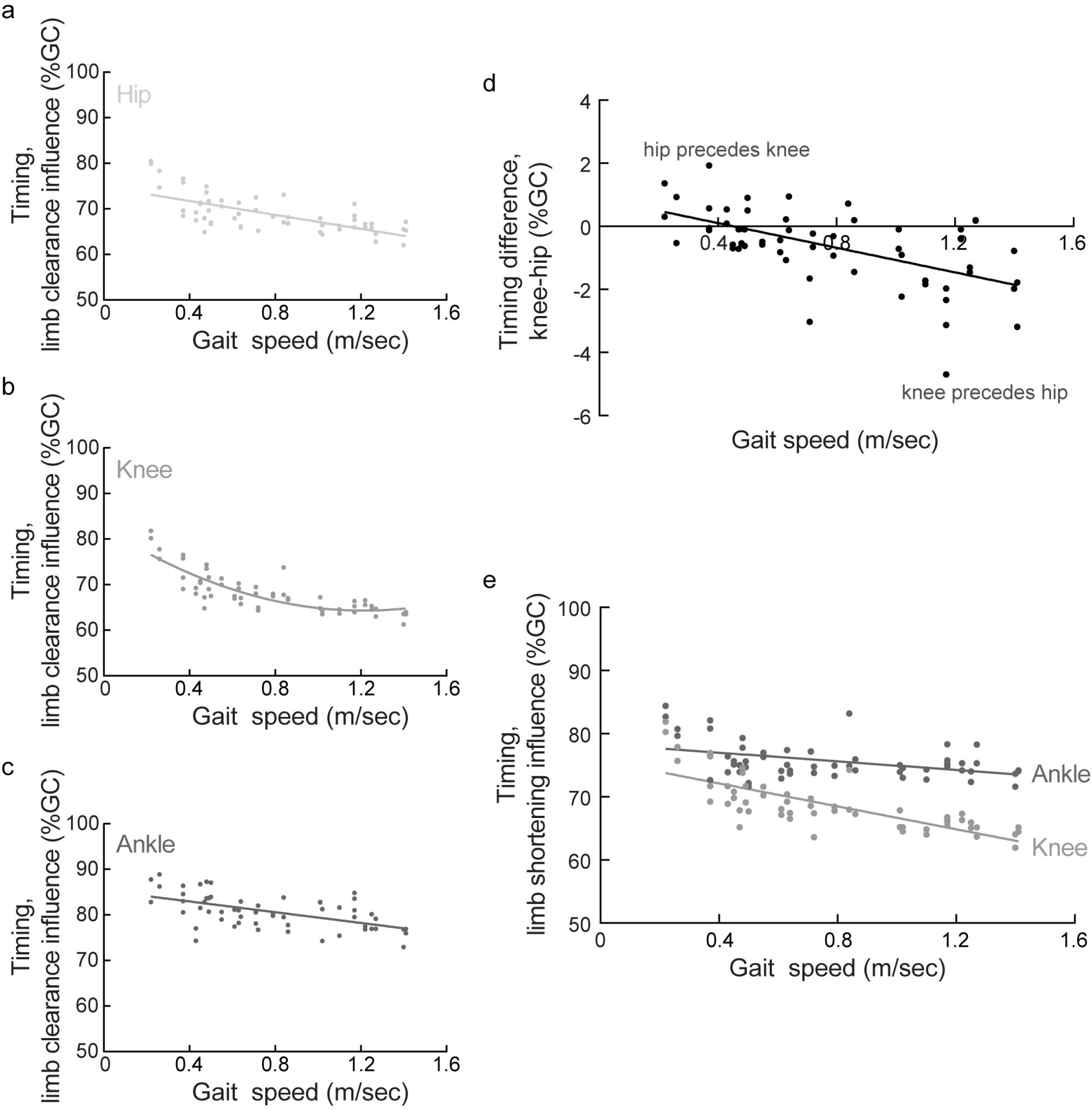
Timing of peak influences with respect to the gait cycle. The timing of peak limb clearance influences represented across gait speeds for the a) hip, b) knee, and c) ankle. Note, for limb clearance the peak knee influence immediately precedes the peak hip influence at typical walking speeds; this temporal pattern reverses with very slow speeds. d) The temporal difference between peak influence from the knee and hip illustrates a reversal in the temporal order of knee and hip influences during very slow walking. e) The timing of peak limb shortening influences represented across gait speeds. The peak knee influence on limb shortening occurs later in the gait cycle at slower walking speeds. The timing of ankle influence remains consistent across walking speeds for limb shortening

The final model for the timing of joint influence on limb clearance revealed that the fixed effects of joint (F_(2,18.6)_=199.7, *p*<0.0001), gait speed (F_(1,19.8)_=93.5, *p*<0.0001), and a quadratic term for gait speed (F**(1,115.8)**=26.2; *p*<0.0001) were significant predictors of the timing of limb clearance influences. Further the intercepts were noted to vary by joint. At typical walking speeds, the peak influence from the knee immediately precedes the peak hip influence, followed considerably later by the peak ankle influence (Figure 5a). Individual joint models revealed the fixed effect of gait speed was a significant predictor of the timing of joint influence for the hip (*b*=−7.76; *p*<0.0001; Figure 5a), knee (*b*=−10.91; *p*<0.0001; Figure 5b), and ankle (*b*=−5.77; *p*=0.0006; Figure 5c), respectively. Additionally, the quadratic term for gait speed remained significant for the timing of knee influence on limb clearance (*b*=12.14; *p*=0.0089). While still tightly coupled, our exploratory analysis revealed a reversal of the temporal order of the peak hip and knee influences that occurs at walking speeds ≤0.45 m/sec (F_(1,19.0)_=16.5; *p*=0.0007; Figure 5d).

Regarding the timing of joint influence on limb shortening, the final model revealed that the fixed effects of joint (F_(1,94.8)_=255.2; *p*<0.0001), gait speed (F_(1,18.0)_=50.8; *p*<0.0001), the interaction of joint x gait speed (F_(1,16.1)_=9.3; *p*=0.0077), and a quadratic term for gait speed (F_(1,117.1)_; *p*<0.0001) were significant predictors. Individual joint models revealed the fixed effects of gait speed was only a significant predictor of limb shortening influence timing for the knee (F_(1,18.8)_=24.9; *p*<0.0001), yet gait speed was not a significant predictor of the timing of influence from the ankle (F_(1,14.0)_=1.9; *p*=0.1844). The relationship of the timing of knee influence on limb shortening reveals a moderate negative slope (*b*_knee_=−7.94; *p*<0.0001). While the changes noted at the knee across gait speeds is considerable, the timing of ankle influence on limb shortening does not change across gait speeds (*b*_ankle_=−2.14; *p*=0.1844; Figure 5e).

## Discussion

### Overview

We note changes in both temporal coordination and the relative joint contributions for limb clearance and limb shortening during very slow walking. Regarding limb clearance, the temporal coordination between the hip and knee changed at very slow walking speeds and the magnitudes of joint contribution reduced with slower walking speeds. While the joint contributions from the hip and knee reduced considerably from the hip and knee, the change in ankle contribution was more subtle. This combination of changes revealed similar contributions from the hip and ankle at very slow walking speeds with motion from the knee still contributing the dominant influence over limb clearance. For limb shortening, the knee contribution was both reduced in magnitude and occurred later in the gait cycle with very slow walking speeds. In contrast, the magnitude and timing of the ankle contribution remained consistent across gait speeds.

### Methodological context

Prior work has used experimental manipulation of walking speeds within a sample to demonstrate the relationship between gait characteristics and gait speed.(Nymark et al. 2005; Oberg, Karsznia, and Oberg 1994; Stoquart, Detrembleur, and Lejeune 2008) Here we studied a single group of individuals walking at two-to-three progressively slower self-paced speeds to investigate coordination over a range of walking speeds. By allowing participants to determine their progressively slower speeds, they were able to self-organize their walking behavior according to their individual biomechanical constraints.

At slow walking speeds, the local minimum of the toe marker trajectory that typically characterizes minimal toe clearance (MTC)(Moosabhoy and Gard 2006; Murray and Clarkson 1966; Begg et al. 2007) is often absent.(Santhiranayagam et al. 2017) Indeed, in our sample the local minimum of limb clearance observed in at comfortable walking speeds was absent during very slow walking. Therefore, we investigated peaks of influence serving limb clearance and shortening, rather than investigating influence at a given gait event. As a result, we captured the relationship between the maximal contributions from the hip, knee and ankle for limb clearance and the knee and ankle for limb shortening. Motion about the knee, specifically knee flexion, was the predominant influence for both limb clearance and limb shortening.

The influence values investigated here were derived from kinematic data. The magnitude of kinematic excursions is known to vary with gait speed; i.e., smaller joint motions are produced with slower walking speeds.(Kirtley, Whittle, and Jefferson 1985; Nymark et al. 2005; Oberg, Karsznia, and Oberg 1994; Stoquart, Detrembleur, and Lejeune 2008; Mentiplay et al. 2018) Yet, the changes in joint contributions cannot be attributed solely to reductions in gait speed given the unchanging ankle contribution to limb shortening.

### Pattern of joint contributions to limb clearance and limb shortening

Visual inspection of the influence curves (Figure 3a) reveals the knee influence over limb clearance is primarily positive throughout swing; in contrast, the hip begins by reducing limb clearance early in swing and then contributes to increased limb clearance towards the end of swing. The absolute magnitudes of these influences decrease with slower walking speeds (Figure 3b). Similarly, the knee influence over limb shortening (Figure 4) has a significant positive contribution (negative peak) early in swing that decreases with slower walking speeds. Importantly, our results reveal a small ankle influence for limb shortening independent of walking speed.

Consistent with prior work, our results demonstrate that knee flexion is the primary sagittal plane driver of limb clearance while ankle dorsiflexion only mildly affects limb clearance.(D. A. Winter 1992) However, any joint motion in the stance or swing limbs, and the muscles that control those motions, can contribute to limb clearance. As an example of this critical point that limb clearance results from a multitude of factors, frontal plane pelvic motion, specifically contralateral hip abduction, has been shown to be highly influential on limb clearance.(D. A. Winter 1992) Our current model accounted for only the sagittal plane motion of the swing limb, thus cannot provide a complete picture of the three-dimensional and contralateral contributions to limb clearance. However, our findings do reveal a clear indication of the relationship between swing limb joint motions and limb clearance and limb shortening.

### Is slow walking a different behavior?

Prior work has noted the relationships between gait speed and certain spatiotemporal characteristics of gait differ at walking speeds <0.5 m/sec. Observation of these changes in characteristics of the walking pattern at speeds below 0.5 m/sec led to the suggestion that very slow walking may constitute a different motor program.(Leiper and Craik 1991; Smith and Lemaire 2018) Here we investigated whether walking at very slow speed simply involves decreased magnitude of the walking pattern or altered coordination thus suggesting a distinctly different gait pattern.

Temporal coordination of the walking pattern changed with very slow walking. Similar to other reports, we found the peak influences on limb clearance from the hip and knee are tightly coupled under all walking conditions with the knee influence immediately preceding the hip influence.(Hershler and Milner 1980; Charteris 1982; Leroux, Fung, and Barbeau 1999; Awai and Curt 2014) However, the temporal order of hip and knee influence reverses at gait speeds < 0.45 m/sec. Thus, the temporal characteristics of very slow walking appear to be altered compared to walking at comfortable speeds. These findings are consistent with previous work that has posited that self-selected and slow walking may be different behaviors.(Leiper and Craik 1991)

Influence values, as used here, represent more than the magnitude of joint excursion. Influence represents the relational contribution of the joint excursion to the functional tasks of limb clearance and limb shortening(Little, McGuirk, and Patten 2014), that is the concurrent coordination between joints to achieve a task goal. Furthermore, influence quantifies the respective joint contributions that are temporally linked to their task goal throughout the gait cycle. We observed significant reductions in the magnitude of hip and knee joint influence with slower walking speeds. Despite these changes, the knee remains the primary contributor to limb clearance and limb shortening at very slow walking speeds. The changes in the magnitude of hip influence bring the hip and ankle contributions into an overlapping range, thus the ankle appears to take on a relatively greater role in limb clearance at very slow speeds. However, the ankle contribution to limb clearance is still reduced at very slow walking speeds compared to comfortable walking speeds.

Although we observed notable, systematic changes in the hip and knee contributions, the ankle contribution changed only minimally across walking speeds. These findings are consistent with the significant body of work illustrating the nervous system controls the endpoint of the limb trajectory whereby motion of distal limb segments are more invariant than proximal segments.(Ivanenko et al. 2002a, 2002b, 2007, 2008; Bosco, Poppele, and Eian 2000) Endpoint control explains the remarkable constancy noted in the relationship between limb velocity and endpoint trajectory across speeds ranging 0.19 – 1.39 m/s.(Ivanenko et al. 2002b) This range of speeds is generally consistent with the speeds produced in our study. Given the premise of endpoint control, the mostly consistent ankle influence we noted across walking conditions for limb clearance and limb shortening is not surprising. Our data illustrate the magnitude of hip and knee influences adjust in order to maintain an invariant contribution from the ankle.

### Significant implications for aging

Our findings also reveal significant implications regarding age-related changes in gait. Not only do older adults walk slower than their younger counterparts, but alterations in intralimb coordination with aging have also been described.(Noce Kirkwood et al. 2018; Byrne et al. 2002; D. A. Winter 1991) The changes in coordination we noted during very slow walking, coupled with the coordination changes attributable to aging, may combine significantly increasing the challenge of slow walking to the point it is unachievable for some. This premise was advanced by Leiper and Craik following observation that the ability of older adults to modulate walking to very slow speeds was systematically related to physical activity levels(Leiper and Craik 1991). Further investigation of this phenomenon is indicated.

### Is speed-matching in biomechanical analyses appropriate?

Speed-matching is commonly used to provide appropriately-scaled normative values when comparing gait parameters between individuals with pathology and non-disabled controls.(Chen et al. 2005) Of note, the slowest gait speeds studied here were exceptionally slow speeds for healthy individuals, but match well with gait speeds observed in clinical populations.(Olney, Griffin, and McBride 1994; Chen and Patten 2008) Observation of altered coordination patterns with very slow walking, which suggest an inherently different motor behavior than walking at comfortable speeds, motivates careful consideration of the outcome measures used for the study of pathologic gait to determine appropriateness of speed-matching. Kinematic measures and certain spatiotemporal variables tend to maintain a consistent relationship with gait speeds, even at very slow walking speeds.(Nymark et al. 2005; Stoquart, Detrembleur, and Lejeune 2008; Mentiplay et al. 2018) For assessment of these metrics, speed-matching is both appropriate and necessary to differentiate persistent gait deviations attributable to pathology rather than walking speed alone.(Jonkers, Delp, and Patten 2009; Chen et al. 2005) On the other hand, investigation of coordination is less straightforward. Given accumulating evidence that very slow walking may be a different motor behavior, it remains to be seen whether coordination patterns adapt and the time required to achieve stable, steady-state performance. These are potential areas for future study.

### Conclusion

Our data illustrate that the temporal coordination and the relative joint contributions serving limb clearance and limb shortening change at very slow walking speeds. These changes support the premise that very slow walking could be a unique behavior, compared to walking at comfortable speeds. Our findings provide new insight regarding temporal coordination at gait speeds below 0.45 m/sec and have significant implications for the study of pathologic gait.

## Disclosure of potential conflicts of Interest

The authors declare that they have no conflict of interest.

## Research involving Human Participants

All procedures performed in studies involving human participants were in accordance with the ethical standards of the institutional and/or national research committee and with the 1964 Helsinki declaration and its later amendments or comparable ethical standards.

## Informed consent

Informed consent was obtained from all individual participants included in the study.

## Acknowledgements

This research was supported by the Department of Veterans Affairs, Rehabilitation Research & Development Service Research Career Scientist Award #N9274S (CP) and NIH T32 Neuromuscular Plasticity Training Grant (VLL; No. 5 T32 HD043730-08, K Vandenborne, PI).

This material is the result of work supported with resources and the use of facilities at the NF/SG Veterans Administration Health Care System, Gainesville, FL, USA and the VA Palo Alto Health Care System, Palo Alto, CA, USA. The contents do not represent the views of the Department of Veterans Affairs or the United States Government. The funding source played no role in either writing this manuscript or the decision to submit for publication. The corresponding author retains full access to all data in the study and assumes final responsibility for the decision to submit for publication.

We thank the following individuals: Ilse Jonkers, PhD for conduct of patient assessments; Eric L. Topp, MS for mathematical review; Drs. Sam Wu, BJ Fregly, and Mark Bishop for helpful comments on an early version of the manuscript, and the study participants for their time and cooperation.

A portion of this work has been presented in abstract form at the annual meeting for the American Society for Biomechanics, August 2012 and Combined Sections Meeting (APTA), January 2013. This work was conducted in partial fulfillment of the requirements for the Ph.D. degree by Virginia L. Little.

